# Next generation protein-based materials capture and preserve projectiles from supersonic impacts

**DOI:** 10.1101/2022.11.29.518433

**Authors:** Jack A. Doolan, Luke S. Alesbrook, Karen B. Baker, Ian R. Brown, George T. Williams, Jennifer R. Hiscock, Benjamin T. Goult

## Abstract

Extreme energy dissipating materials are essential for a range of applications. The military and police force require ballistic armour to ensure the safety of their personnel, while the aerospace industry requires materials that enable the capture, preservation and study of hypervelocity projectiles. However, current industry standards display at least one inherent limitation. To resolve these limitations we have turned to nature, utilising proteins that have evolved over millennia to enable effective energy dissipation. Specifically, a recombinant form of the mechanosensitive protein talin was incorporated into a monomeric unit and crosslinked, resulting in the production of the first reported example of a talin shock absorbing material (TSAM). When subjected to 1.5 km/s supersonic shots, TSAMs were shown not only to absorb the impact, but to capture/preserve the projectile, making TSAMs the first reported protein material to achieve this.

## Main

When impacted by a projectile, a material is exposed to a variety of phenomena simultaneously. To survive the impact a material must contend with wave propagation (elastic, shock and plastic), fragmentation, perforation and spallation^1^. Thus, installing a mechanism within a material to enable effective energy dissipation is essential for multiple applications^2-4^. Body armour is commonly used by military and civilian forces to protect the wearer against penetration from projectiles, such as bullets or shrapnel^4^. Frequently, this armour consists of a multi-layered system, commonly a ceramic face backed by a fibre-reinforced composite^5^. This multi-layered design enables the hard brittle ceramic to destroy the projectile tip, in turn distributing the kinetic energy over the backing which reflects the tensile wave and captures the shattered ceramic^6^. Despite the effective penetration blocking of these armour systems, a remainder of the kinetic energy is still distributed to the wearer, often resulting in behind armour blunt trauma^7^. Furthermore, during impacts this form of armour is irreversibly damaged, compromising its structural integrity for further use. The aerospace sector utilise impact energy dissipating materials for the unique task of capture and preservation of space debris, space dust and micrometeoroids^8^. These captured projectiles contribute towards our understanding of the local environments of aerospace equipment, including that of the international space station^9^. Data from these experiments facilitate aerospace equipment design, improving the safety of astronauts and the longevity of costly aerospace equipment. Aerogels are the current industry standard for projectile capture and preservation, achieving energy dissipation through conversion of projectile kinetic energy into both mechanical and thermal energy^10^. However, the resulting temperature elevation, further enhanced by the remarkable insulating properties of aerogel^11^, can cause the aerogel structure to melt^10^. Furthermore, these elevated temperatures may compromise the structure of the captured projectiles, altering its chemical composition^10, 12^. This thermal and mechanical energy, causes chemical bond breakage, rendering the aerogel irreversibly damaged post-impact. It is apparent from the aforementioned examples that a material utilising an energy dissipation mechanism that reforms following the removal of force would alleviate inherent issues seen with the industry standard materials. Additionally, specifically for the aerospace sector, energy dissipation that does not result in the conversion of kinetic to thermal energy would be beneficial.

Within the animal kingdom, proteins that offer unique mechanical properties are rife; silk fibroin displays modifiable macroscale properties in its assembled fibre form, while elastin instils elasticity in animal tissues^13^. Although there are many proteins analogous to these examples, very few researchers have tapped into these natural resources for development of materials with novel mechanical properties^14-16^ ; even fewer have tested these materials for real world applications outside of the biomedical sector^13^. Talin (Fig. 1a) is the epitome of a mechanical protein, mediating the connection between the actin cytoskeleton and the integrin extracellular matrix receptors, acting as a mechanosensor. Previous work determined that, through unfolding/refolding events of its thirteen four/five helical rod domains^17, 18^,^19, 20^ when stretched within the physiologically relevant range, talin is able to maintain the average force experienced by the protein below 10 pN^18^. Furthermore, upon removal of force, refolding of the talin rod domains occur with high fidelity over numerous force cycles^18^ confirming talin as a cellular shock absorber.

**Fig. 1.**
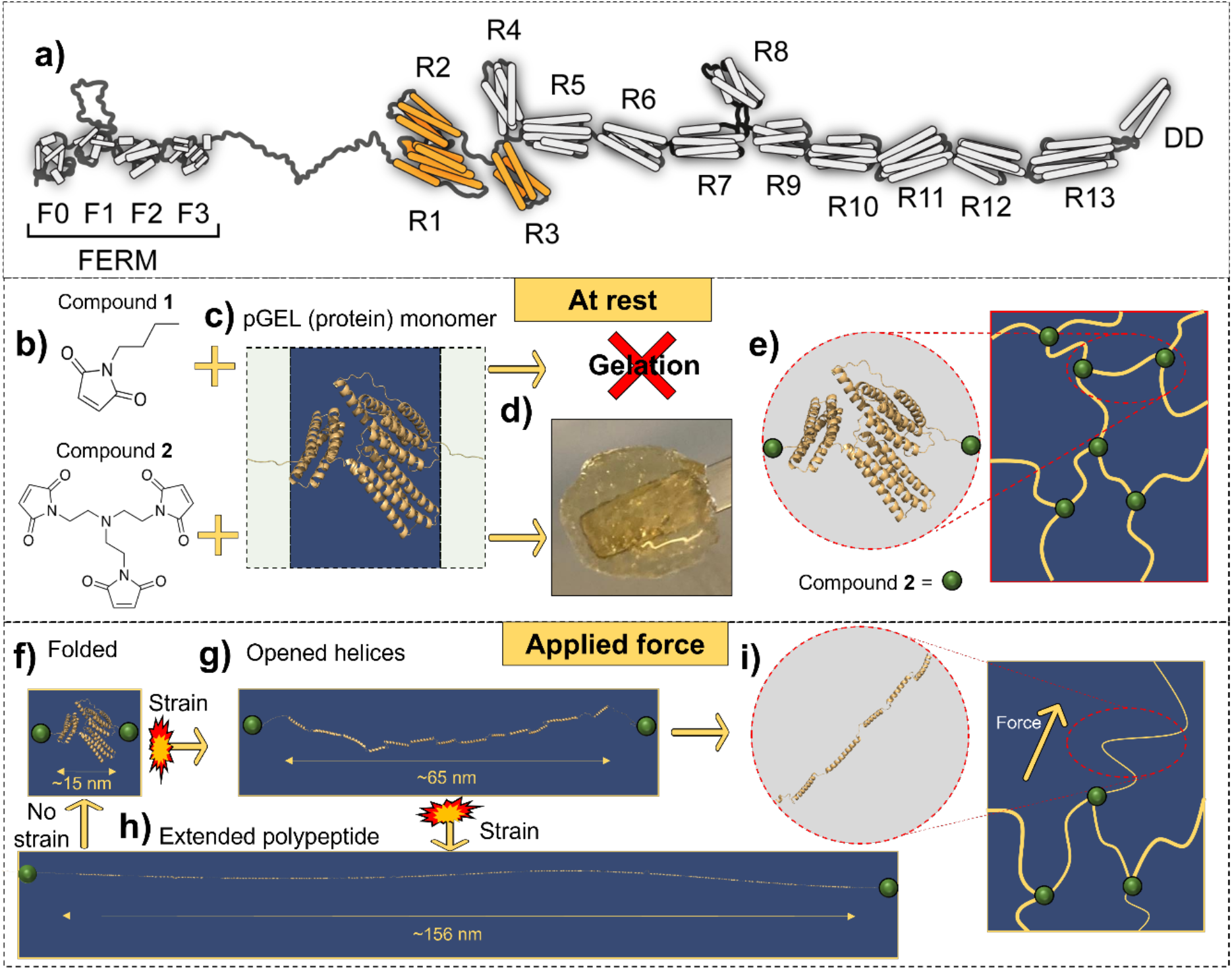
The design concept of TSAM. **a**. Cartoon representation of the protein talin, F = FERM domain, R = rod domain, DD = dimerisation domain. The R1-R3 region is shown in orange. **b**. The control monovalent crosslinker **1** (Fig. S2 and S4) and the trivalent crosslinker **2** (Fig. S3-S4), **c**. pGEL in the folded state, green boxes = flexible linkers, blue box = R1-R3 domains of talin. **d**. Resulting gelation for each crosslinker. **e**. Representation of the three-armed network structure formed from crosslinker **2** with no applied strain. **f**. pGEL in fully folded state presents length of ≈15 nm. **g**. When exposed to strain pGEL unfolds into a linear string of helices extending to ∼65 nm in length. **h**. When exposed to higher strain, pGEL unfolds fully into extended polypeptide, increasing to a length of ∼156 nm. Complete refolding can occur once strain is removed. **i**. Representation of the three-armed network structure with applied strain, causing extension of protein into opened helices form, increasing fibre length^22^.

Here, we have engineered a recombinant form of talin, termed pGEL, which comprises three rod domains of talin, R1-R3, that are modified (with internal cysteine residues mutated to serine and cysteine residues introduced at either end of the protein) for use as the monomer with which to form a polymer. When exposed to force, these three domains provide a stepwise unfolding, with the wild type domains exhibiting threshold unfolding forces of 20, 15 and 5 pN respectively^18^. We hypothesised that, upon application of force (i.e. shear strain or impact), the three rod domains within each protein monomer would unfold, dissipating energy through the endothermic process of protein unfolding^21^ (Fig. 1f-i).

Using compounds **1** (control monovalent crosslinker) and **2** (trivalent crosslinker) (Fig. 1b, Supplementary Fig. 1-3), pGEL (Fig. 1c) was formed into a hydrogel (Fig. 1d-e) via tri-substitution of the terminal cysteines with crosslinker **2** (Supplementary Fig. 4). The resulting hydrogel, which we have termed TSAM (Talin Shock Absorbing Material), therefore contains monomeric units capable of refolding upon removal of force, retaining its energy dissipating mechanism following any potential impact events. Due to the endothermic energy dissipating mechanisms of protein unfolding^21^ in TSAM, the heating of the captured projectiles seen in aerogels would not be observed, offering a solution to several of the limitations seen with current state of the art impact absorption materials.

### TSAM structural characterisation

The R1-R3 domains of talin incorporated in the pGEL monomer were confirmed to retain alpha helical folding, using circular dichroism and ^1^H-^15^ N HSQC nuclear magnetic resonance (Supplementary Fig. 5-8). Following formation of TSAM, characterisation of the internal network structure was conducted. His-tagged gold immunostaining of the TSAM, imaged using transmission electron microscopy (TEM), confirmed the presence of pGEL in a lattice formation, displaying pore sizes of approximately 100 nm (Fig. 2a). Following this, scanning electron microscopy (SEM) revealed TSAM to contain a porous like structure on the micrometre scale typical of hydrogels (Fig. 2b), with long fibres of width ≈2 µm and pores of ≈10 µm. Elemental dispersive X-ray (EDX) analysis confirmed the observed fibres in the SEM images consisted of sulphur and carbon (Fig. 2c), pGEL representing the only component of the xerogel containing these atoms. Together these findings indicated pGEL molecules linked with crosslinker **2** form a lattice on the nanometre scale, morphing into larger fibrillar like structures on the micrometre scale. When handling TSAMs high levels of extensibility were observed, presenting extension of >3-fold when under tension, and returning to original size upon removal of force (Fig. 2d-e).

**Fig. 2.**
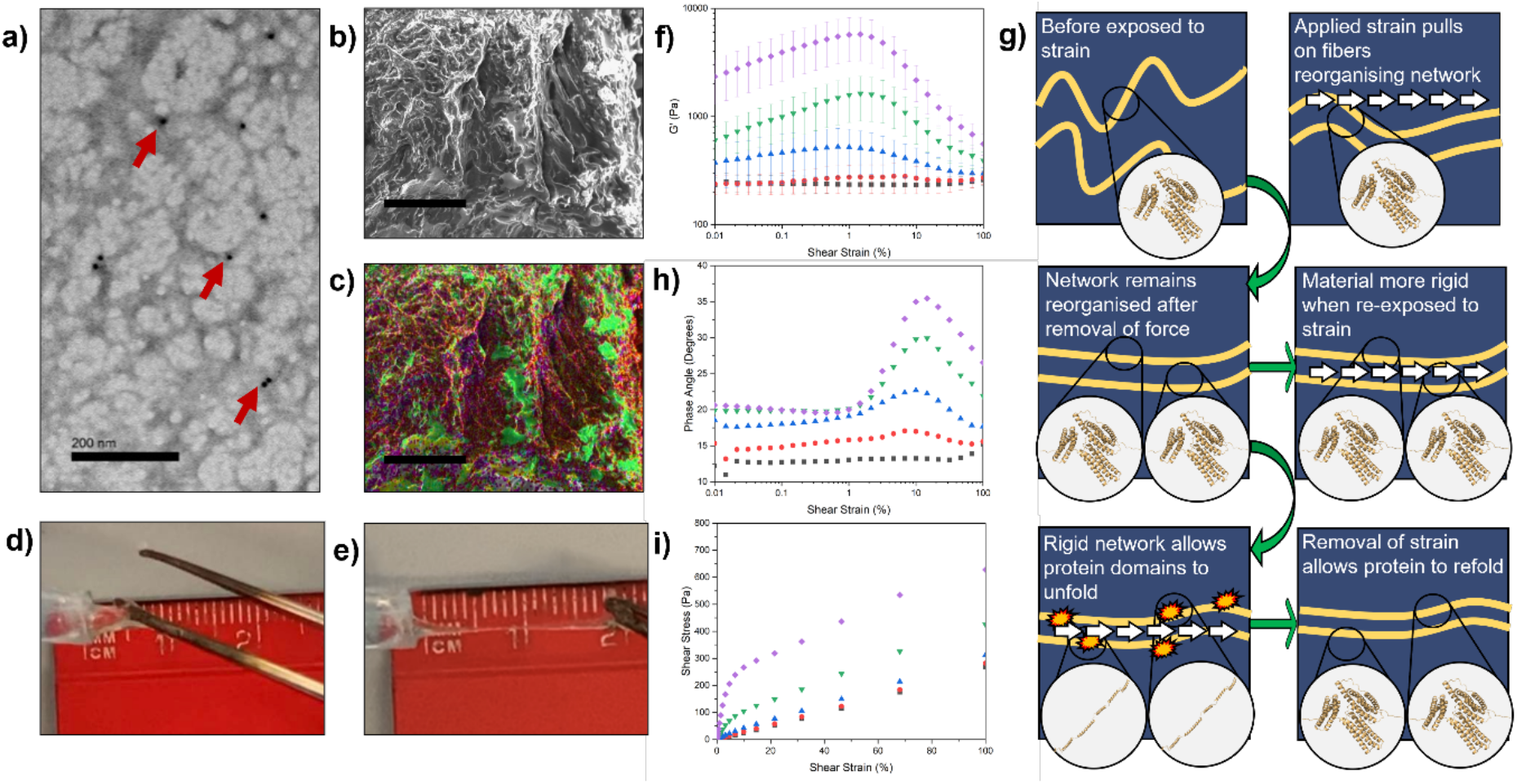
The internal fibre structure of TSAM and its macroscale characterization. **a**. Immunogold-stained TSAM imaged with TEM showing lattice structure of connected pGEL proteins. Gold particles are observed as black dots, some of which are highlighted with red arrows (Scale bar = 200 nm). **b**. The dense fibre structure of TSAM displaying a porous network imaged with SEM on secondary electron mode (Scale bar = 50 µm). Pore sizes are on the range of a few µm. **c**. EDX analysis of SEM image in b. sulphur = yellow, carbon = red, oxygen = green, sodium = teal, phosphorus = purple. (Scale bar = 50 µm) **d**. TSAM slightly stretched. **e**. TSAM stretched to 3x its length. **f**-**i**. Rheological measurements of TSAM (n = 3). **f**. G’ as a product of shear strain (error bars = SEM) for sweeps 1 (squares), 2 (circles), 3 (triangles), 4 (inverted triangles) and 5 (diamonds). **g**. Schematic summary of the events hypothesised to occur over 5 x repeated oscillatory sweeps. **h**. Phase angle against shear strain for sweeps 1-5 on TSAM. **i**. Shear stress against shear strain for sweeps 1-5 on TSAM.

### Evidence for pGEL domain unfolding in TSAM

Subsequent rheological characterisation of TSAMs provided strong evidence for the induced unfolding of the talin domains within the material when exposed to shear strain, indicating that the energy dissipating mechanisms of talin were successfully incorporated into TSAMs. Specifically, for the first applied oscillatory shear strain sweep on the TSAM, the dynamic shear storage (G’) and loss modulus (G”), as a product of shear strain presented a linear viscoelastic region (LVER) for the full range of shear strain tested (Supplementary Fig. 9). Following a two-minute recovery period, a subsequent sweep was performed (Supplementary Fig. 10), once again followed by a two-minute rest period. This protocol was conducted a total of five times (Supplementary Fig. 9-13, Fig. 2f). Owing to the unfolding and refolding kinetics intrinsic to R1-R3, we predicted viscoelastic properties would be retained upon repeated exposure to shear strain. The TSAMs presented G’ > G” confirming viscoelastic behaviour for all five consecutive oscillatory shear strain sweeps. Moreover, the complex modulus (G*) (sum of G’ and G”) increased concomitantly with accumulated sweeps, indicating TSAM presented increased resistance to deformation upon repeated exposure to shear strain^23^. Strain stiffening as a consequence of fibre reorganisation is a well-documented phenomenon occurring in hydrogels formed from biopolymers^24^, causing the elastic modulus to increase with strain. Strain stiffening presents here as the positive gradient observed in sweeps 3-5 (Fig. 2f). The peak maxima for G’ was reached between 1-5% for sweeps 3-5, shifting to the right and increasing in amplitude for each subsequent sweep. As stated, we propose this phenomenon to be due to strain stiffening, resulting in a tighter, more rigid network structure with a greater number of talin domains arranged in parallel to the axis of the fibres (Fig. 2g). As a consequence of the increased network rigidity, strain can become imparted on the internal structure of the fibres themselves. When a maximum fibre strain is reached, mass chain unfolding of the protein domains occur, causing the positive slope to transition into a negative gradient as a result of the sudden introduction of slack from the extension of the proteins (Fig. 2g and Fig. 1f-h). Due to the sudden introduction of slack, a lag between the controlled shear strain and measured shear stress sine waves occur, presenting as a sudden shift towards more viscous-like behaviour, hence the negative gradient of G’ (Fig. 2f) and decrease in G*. Alongside the observed bell-shaped trend of G’ and G” with accumulating sweeps, an increase in phase angle was observed, also presenting a bell-curve (Fig. 2h). Interestingly this pattern was shifted to higher strains, with the peak of the bell occurring between 10-15% strain, directly correlating with the negative slope of G’. The large quantity of slack (Fig. 2g) introduced into the system following a mass unfolding event allows for increased flow, observed as a drastic phase shift towards 45 degrees, and decreased rigidity, seen as a drop in G*. With increased application of shear strain, this phase shift peak eventually declines as tension within the fibres begin to increase again. Upon removal of shear strain, unfolded proteins may then refold, and the resulting TSAM displays an enhanced rigidity (higher G* at the start of the next sweep) due to fibre network reorganisation. Shear stress vs. shear strain correlations corroborate these results, revealing an exponential increase in shear modulus (G), a measure of rigidity, with accumulated sweeps, further illustrating the strain stiffening within TSAM (Fig. 2i). Furthermore, sweeps 4 and 5 reach shear yield points, beginning to move into viscous stress as seen by the induction of a slope, subsequently transitioning back into a linear gradient indicating the reoccurrence of elastic behaviour. In summary, the linear elastic region at low shear strain is a result of reordering of the network structure and gradual tension accumulating in the fibres, the following curving transition indicates sudden mass unfolding of talin rod domains, and subsequent linear region reports elasticity reoccurring once tension is again applied to the fibres with increasing shear strain.

To confirm that the unfolding of the talin domains within the TSAM is directly responsible for the rheological characteristics/material properties observed, a green fluorescent protein tagged-vinculin domain 1 protein (GFP-VD1) was employed. GFP-VD1 is capable of selectively binding to the unfolded state of each of the rod domains (Fig. 3a, Supplementary Fig. 14), preventing domain refolding and ‘locking’ the extended conformation^25^. Here the GFP-VD1 was introduced into the TSAM pre-amplitude sweep as a 2 mg/mL solution through a material swelling process. The rheological properties of these materials were then elucidated and compared to the results of analogous studies in which the same TSAM material underwent the same material swelling process in a solution of GFP or buffer only. The resulting G’ and G” as a product of shear strain for the three conditions tested are summarised in Supplementary Fig. 15.

**Fig. 3.**
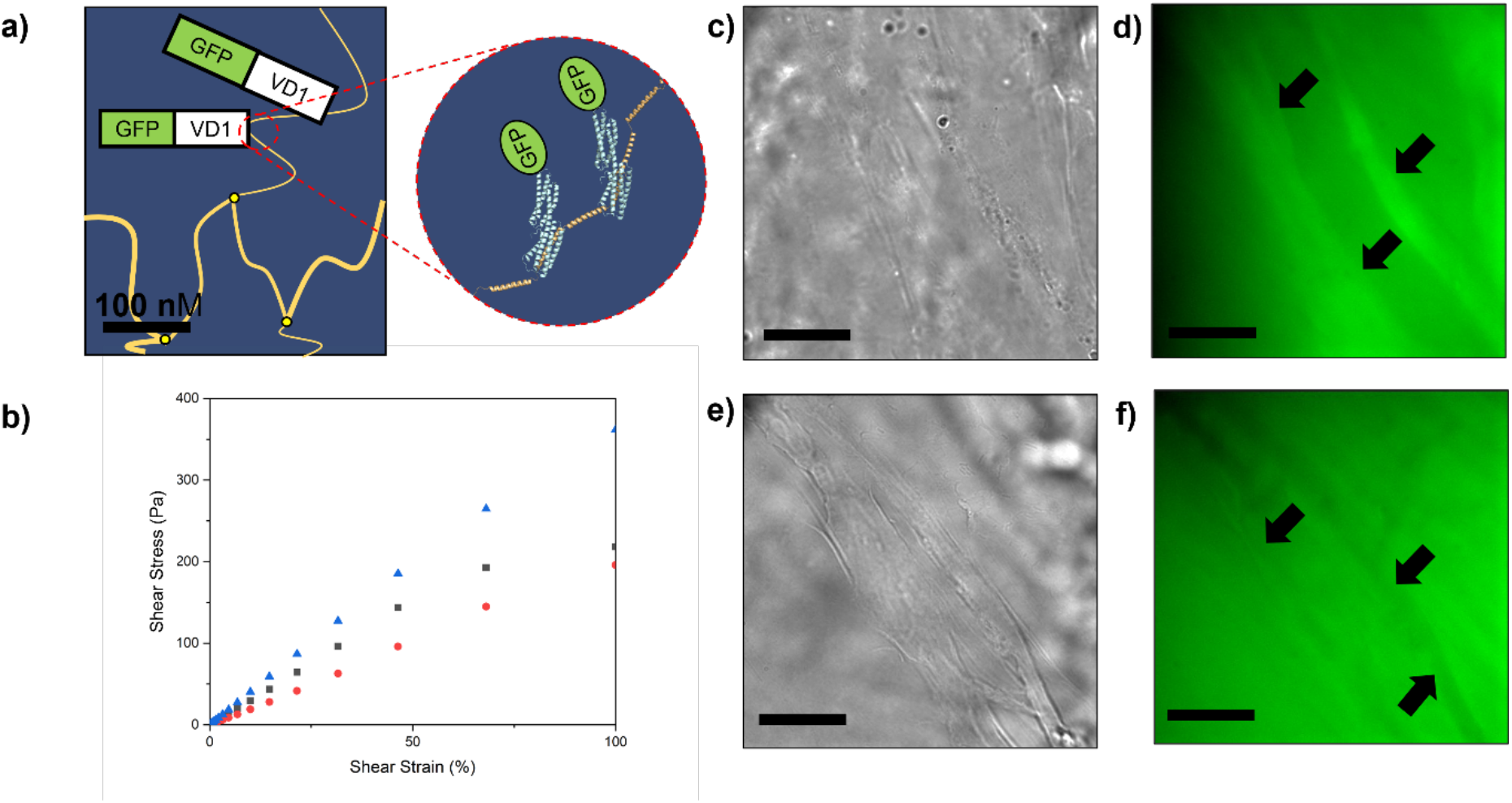
Effects of GFP-VD1 on TSAM. **a**. Representation of GFP-VD1 binding to unfolded pGEL in TSAM fibres, with resulting cartoon protein figures created in PyMOL using VD1 PDB structure 1U6H^26^. **b**. Shear stress as a product of shear strain for buffer (blue triangles), GFP-VD1 (black squares) and GFP (red circles), showing GFP-VD1 treated TSAM reaches its yield point between 46-68% shear strain. **c**. Transmitted light image of GFP-VD1 localised to TSAM fibres (scale bar = 20 µm). **d**. Maximum projection widefield fluorescent image of c. (scale bar = 20 µm) with fibres showing localised GFP-VD1 indicated by arrows. **e**. Transmitted light image of GFP in TSAM (scale bar = 20 µm). **f**. Maximum projection widefield fluorescent image of e. showing GFP sitting in void space, with fibres this time visible as darker structures indicated by arrows (Scale bar = 20 µm).

When plotted as shear stress against shear strain (Fig. 3b) the GFP and buffer controls presented the same linear trend as obtained in the first amplitude sweep for the non-treated TSAMs, indicating purely elastic behaviour. In contrast, the TSAM treated with GFP-VD1 reached a yield point between 46-68% shear strain (Fig. 3b) as a result of VD1 binding events. To further confirm the binding of GFP-VD1 to the TSAM fibres, a series of comparative fluorescence microscopy experiments were conducted. Here, fibre like structures exhibiting the same diameter as those observed in our previous SEM studies (Fig. 2b) were found to have localised GFP-VD1 (Fig. 3c-d), confirming binding. In contrast, the GFP control treated TSAM fibres appeared as darker regions, with void spaces presenting higher GFP concentrations (Fig. 3e-f).

### TSAMs capture and preserve projectiles from supersonic impacts

Following the rheological evidence for TSAMs retention of talin’s endothermic energy dissipation mechanism, we moved on to test the performance of the TSAM as an impact absorbing material, investigating TSAM performance upon supersonic projectile impact. Specifically, velocities of 1.5 km/s were tested, as this is a speed relevant to the aerospace and defence industries.^27, 28^ For instance, particles in space impact both natural and man-made objects at speeds >1 km/s^27^, while muzzle velocities from firearms commonly fall between 0.4-1.0 km/s^28^. Here the TSAM, in addition to a commercially available polyvinylpyrrolidone hydrogel control, were placed in the target chamber of a light gas gun (LGG) and the following material properties elucidated: (1) the ability of the TSAM to survive impact; (2) the ability of the TSAM to reduce the force of the projectile before impacting an aluminium back plate; and (3) the ability of the TSAM to capture the projectile in a preserved state.

Spherical basalt particles between 20-70 µm were used as projectiles, loaded in a sabot as buckshot. A schematic for this experiment is given in Fig. 4a-c. When shot at 1.5 km/s, the control gel was destroyed (Fig. 4d), with a visible hole in the tape behind the gel (Fig. 4e), and a crater of 1.33 mm in diameter produced in the aluminium back plate (Fig. 4f). Therefore, this material control showed no detectable impact absorption properties. However, under analogous experimental conditions, the TSAM appeared mostly intact from the frontal perspective (Fig. 4g), with no projectile permeation detected to either the supporting tape (Fig. 4h) or the aluminium backplate (Fig. 4i). In addition, subsequent SEM analysis identified the basalt particles embedded in the TSAM post shot (Fig. 4j), confirming that the TSAM had completely absorbed the impact of the basalt buckshot. The transparency of the TSAM shown in Fig. 4c and Supplementary Fig. 16 is an additional desirable property, allowing for the easy removal of caught projectiles from the TSAMs. To conclusively determine if TSAM also enabled preservation of the captured basalt projectiles, SEM was performed on the impacted TSAM. Multiple basalt particles presenting a preserved circular shape were observed in the gel (Fig. 4j-k), confirmed as basalt with EDX analysis (Supplementary Fig. 17). Thus, supporting that TSAM enables projectile preservation. Moreover, during one of the TSAM shots, shrapnel from the aluminium (Al 7075) burst disk (Fig. 4b) struck the TSAM in combination with the basalt, as confirmed through SEM and EDX analysis (Fig. 4i and Supplementary Fig. 18). Such an impact often destroys aerogel materials used as the industrial standard within the aerospace industry for projectile capture, providing evidence that TSAMs are able to overcome this limitation.

**Fig. 4.**
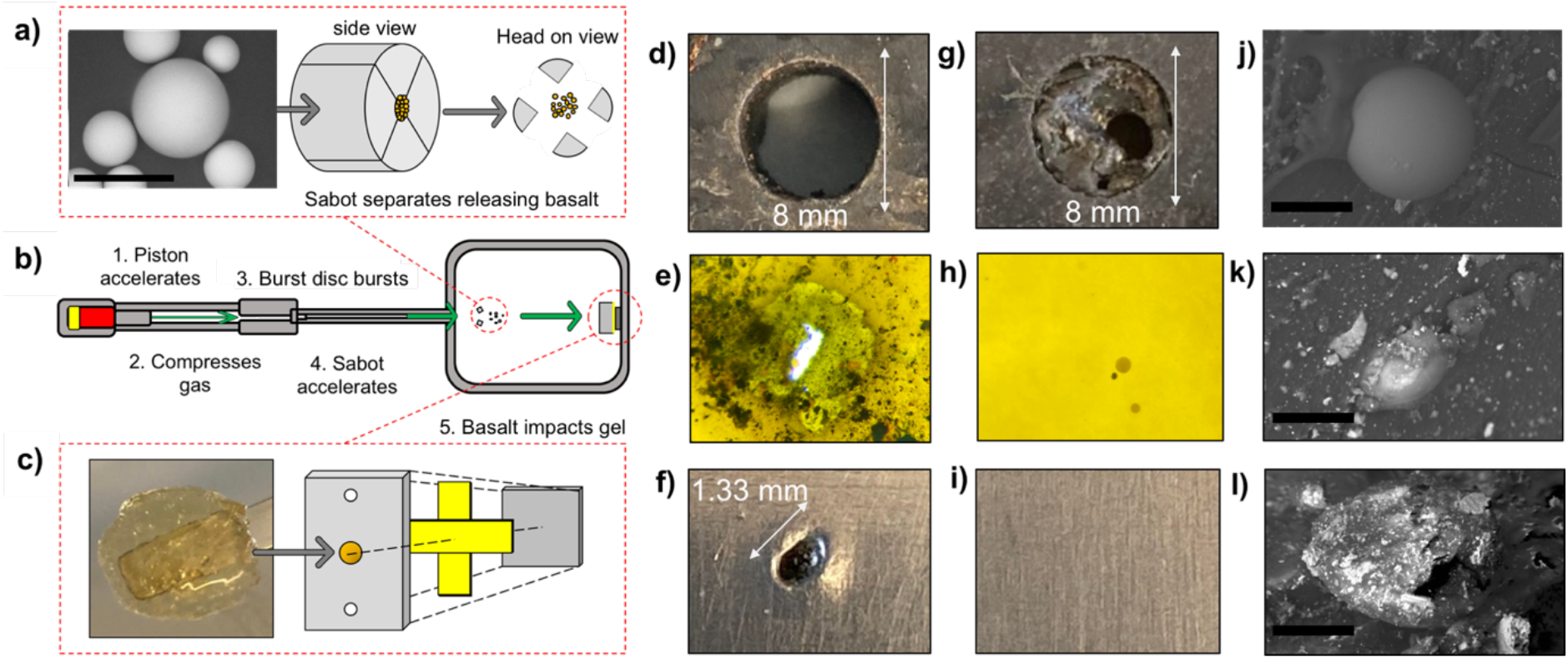
Supersonic impact study on TSAM. **a**. SEM image of a basalt particle used as the projectile and representation of how the basalt is loaded into a sabot and its release during a shot (scale bar = 60 µm). **b**. diagram of the light gas gun apparatus with the key stages after the shot is triggered. **c**. image of TSAM and how it is prepared as a target. The TSAM is loaded into a target plate constructed of steel (Blast tank exit aperture, stainless 304), with tape used to seal the back of the hole, followed by an aluminium back plate (Al 5083). **d**-**f**. results from control gel **d**. Destroyed control gel after basalt impact at 1.5 km/s. **e**. Hole formed in tape from basalt projectile. **f**. crater formed in aluminium back plate. **g**-**I** results from TSAM **g**. mostly intact TSAM after basalt impact at 1.5 km/s. **h**. tape with no hole, containing several caught basalt particles in the transparent TSAM attached to its surface (Supplementary Fig. 16). **i**. undamaged aluminium back plate. **j**. SEM image of intact basalt particle caught by TSAM after impact at 1.5 km/s. (scale bar = 45 µm) **k**. SEM image of another basalt particle caught by TSAM after impact at 1.5 km/s. (scale bar = 30 µm) **l**. SEM image of a fragment of the aluminium (Al 7075) burst disc that impacted TSAM during the 1.5 km/s basalt shot (scale bar = 50 µm).

## Discussion and Outlook

In summary, we present the first example of a talin shock absorbing material (TSAM) known to literature – a SynBio material constructed from monomeric units containing force-dependent mechanical switch domains. In addition, we show that TSAMs can absorb impacts by basalt particles and larger pieces of aluminium shrapnel, providing the first example of a protein material capable of absorbing supersonic projectile impacts. These results lend the TSAMs towards application within the aerospace and defence industries, e.g. as a backing for multi-layered armour where shattered ceramic capture is required, and in hypervelocity impact experiments in which the projectile needs to be preserved for further study. As a consequence of the endothermic energy dissipating mechanism of talin, it is very unlikely any elevation of projectile temperature was induced during the LGG experiment^21^, offering a distinct benefit over aerogel materials. This energy dissipating mechanism was confirmed using rheology while, GFP-VD1 binding experiments supported the presence of talin unfolding events within these processes. Through the reversible refolding of talin domains within TSAM following the removal of force, the material also demonstrates potential for iterative use. Finally, as talin contains thirteen helical domains, each with unique unfolding forces, these TSAMs may be tuneable by modifying the talin domains featured in the monomer unit, offering the potential for tailoring toward a diverse array of mechanical properties and resulting applications.

## Methods

### Protein engineering

The genes encoding pGEL, GFP-VD1 and GFP were constructed in pET151 vectors. The proteins were expressed in BL21(DE3)* *E. coli*. Protein purification was achieved using HisTrap HP columns (Cytiva) for His-tag based affinity chromatography using an AKTA Start protein purification system (Cytiva). Following purification, proteins were dialysed in phosphate buffer (20 mM sodium phosphate, pH 7.4, 50 mM NaCl).

### TSAM preparation

A 30:1 ratio of TCEP:cysteine was slowly added to a solution of pGEL in phosphate buffer (pH 7.4). After one hour, the pGEL solution was run through PD10 desalting columns (Cytiva) twice to ensure TCEP removal. Immediately following the desalting step, the pGEL solution was concentrated to the desired concentration using 30 kDa MWCO concentrators (SigmaAldrich). The TSAM was then formed through the addition of crosslinker 2 at 1:1 maleimide:sulfhydryl molar ratio. The TSAM was left to set at 4°C overnight.

### Scanning electron microscopy

The TSAM was dehydrated to form a xerogel and placed on a carbon tab mounted onto an aluminium stub. Imaging was achieved using a Hitachi S-3400N scanning electron microscope with elemental dispersive X-ray analysis and analysed using Oxford instruments AZtec software.

### Immuno-gold staining and transmission electron microscopy

2 µl of sample was applied to carbon/formvar 400 mesh gold grids (Agar Scientific) and allowed to settle on the grid for 5 minutes. The sample was then fixed in 2% formaldehyde and 0.5% glutaraldehyde in 100 mM sodium cacodylate buffer pH 7.2 (CAB) for 15 minutes at room temperature. Samples were washed 2 × 5 minutes in CAB and 2 × 5 minutes in 20 mM Tris, 500 mM NaCl, 0.1% BSA and 0.5% Tween 20 (TBST). Grids were blocked in 2% BSA in TBST for 30 minutes and then moved into a 20 µl drop of anti-His tag primary antibody (Sigma) diluted 1:100. Grids were washed 6 × 2 minutes in drops of TBST before incubation in Goat anti-mouse IgG conjugated to 5 nm gold particles (British Biocell International) diluted 1:50 for 30 minutes. Grids were washed for 6 × 2 minutes in TBST and 6 × 2 minutes in distilled water. Negative controls were performed as above but primary antibody was replaced with TBST. Samples were then air dried and negative stained in 2% aqueous uranyl acetate. Samples were viewed using a Jeol 1230 Transmission electron microscope at 80 kV and images were recorded on a Gatan OneView 16 MP digital camera.

### Rheological measurements

Rheological measurements were performed on an Anton Parr modular compact rheometer (MCR302). All measurements were performed at 298 K using a PP20 parallel plate. Oscillatory amplitude experiments maintained a frequency of 10 rad/sec and were performed with an amplitude of oscillation range of 0.01-100%. A 2 minute rest time was set between each amplitude sweep, with a total of five sweeps performed on each TSAM. For the GFP-VD1, GFP and buffer swelled experiments, the TSAM was left in 2 mg/mL of the respective solution overnight before rheological measurements were performed.

### Fluorescence microscopy

GFP-VD1 and GFP treated samples of TSAM from the rheology experiments were visualised using an Olympus IX71 microscope employing a 1.6x magnification Optovar in combination with a PlanApo 100x OTIRFM-SP 1.49 NA lens mounted on a PIFOC z-axis focus drive (Physik Instrumente, Karlsruhe, Germany), and illuminated using LED light sources (Cairn Research Ltd, Faversham, UK) with DC/ET350/50x excitation, ET Quad Sedat dichroic, and DC/457/50m emission filters (Chroma, Bellows Falls, VT). Samples were visualised using a QuantEM (Photometrics) EMCCD camera, and the system was controlled with Metamorph software (Molecular Devices). Each 3D-maximum projection of volume data was calculated from 31 z-plane images and the best 6 were chosen, each 0.2 µm apart, and analysed using MetaMorph software.

### Light gas gun experiments

The impact experiments were carried out using the Light Gas Gun (LGG) facility at the University of Kent, Canterbury. The LGG is capable of accelerating projectiles smaller than 3.5 mm to speeds up to 7 km/s^29, 30^. The TSAM target was set in a blast tank exit aperture (BTEA) with a circular, 8 mm diameter aperture, sealed with tape, with an aluminium (5083) back plate placed behind. Multiple 20-70 °m basalt particles were loaded into a single sabot utilising the “buckshot” method and were fired at roughly 1.5 km/s, with the speeds recorded via the BTEA - Muzzle laser method as described by Burchell et al.^29^. The target was removed prior to the air flushing procedure to reduce gun contamination of the TSAM. The combination of the BTEA and target mount into a single device, allowed for minimal spreading of the buckshot projectile, increasing the chance of direct impact onto the TSAM, and maximized the BTEA-muzzle separation.

## Acknowledgements

BTG would like to acknowledge BBSRC (BB/S007245/1), Cancer Research UK Program grant (DRCRPG-May21) for funding and The Royal Society Project grant (RGS\R2\192016). JAD would like to acknowledge the University of Kent for funding. JRH would like to thank the UKRI for the funding of her Future Leaders Fellowship (MR/T020415/1).

The TSAM material is pending a UK Patent Application No. GB2216633.4

