## Extended Supplementary Information for "Next generation protein-based materials capture and preserve projectiles from supersonic impacts"

#### Table of Contents

***Section 1: Supplementary methods*** ... **2**

***Section 2: Chemical structures*** ... **3**

Supplementary Fig. 1 ... 3

***Section 3: Chemical synthesis and characterisation*** ... **4**

**Chemical synthesis:** ... **4**

**NMR characterisation:** ... **5**

Supplementary Fig. 2 ... 5

Supplementary Fig. 3 ... 5

**Electrospray ionisation mass spectrometry of crosslinkers 1 and 2 forming substituted species with Kank1 KN-domain peptides.** ... **6**

Supplementary Fig. 4 ... 6

***Section 4: pGEL characterisation*** ... **7**

**Circular Dichroism confirming preserved folding between pGEL and wild type R1-R3.** ... **7**

Supplementary Fig. 5 ... 7

**Circular Dichroism confirming preserved folding between wild type R2 and the isolated mutated R2 domain from pGEL.** ... **7**

Supplementary Fig. 6 ... 7

**2D HSQC spectra of $^{15}$N-labelled mutated R1 showing folding is retained compared to the wild type R1.** ... **8**

Supplementary Fig. 7 ... 8

Supplementary Fig. 8 ... 9

***Section 6: Rheology of TSAM*** ... **9**

**Amplitude Sweep 1** ... **9**

Supplementary Fig. 9 ... 9

**Amplitude Sweep 2** ... **9**

Supplementary Fig. 10 ... 9

**Amplitude Sweep 3** ... **9**

Supplementary Fig. 11 ... 9

**Amplitude Sweep 4** ... **10**

Supplementary Fig. 12 ... 10

**Amplitude Sweep 5** ... **10**

Supplementary Fig. 13 ... 10

***Section 7: TSAM with GFP-VD1, GFP and buffer rheology measurements*** ... **11**

Gel filtration confirming GFP-VD1 binds to pGEL. 11
Supplementary Fig. 14 11

Amplitude sweep 1 11
Supplementary Fig. 15 11

*Section 8: Light gas gun (LGG) experiments* 12

Supporting images from LGG experiments 12
Supplementary Fig. 20 12

EDX analysis of caught basalt particles from light gas gun experiments imaged in SEM. 13
Supplementary Fig. 21 13

EDX analysis of caught burst disc shrapnel in light gas gun experiment imaged with SEM. 14
Supplementary Fig. 22 14

*References* 15

#### Section 1: Supplementary methods

**Compound NMR characterisation:** NMR spectra for compounds **1** and **2** were obtained on a Bruker AV2 400 MHz spectrometer. The data was processed using TopSpin. NMR chemical shift values are reported in parts per million (ppm) and calibrated to the centre of the residual solvent peak set (s = singlet, br = broad, d = doublet, t = triplet, q = quartet, m = multiplet).

**Electrospray Ionisation mass spectrometry:** ESI-MS was performed on an Agilent HPLC system connected to a Bruker micrOTOF-Q mass spectrum instrument. Spectra were analysed using Bruker's Compass Data Analysis software. All samples were run using solvent A (0.05% TFA in water) and solvent B (80% acetonitrile, 0.045% TFA in water). Samples were prepared at a concentration of 100 μM peptide in phosphate buffer (20 mM $NaH_2PO_4.2H_2O$, 50 mM NaCl, pH 7.4) and reduced with 5 mM TCEP. Following a 10 minute reduction time the respective compound was added at a 10:1 ratio and allowed to react for two hours. A total of 5 μL of the sample was then loaded into the LCMS.

**pGEL01 sequence design:**

MHHHHHHGKPIPNPLLGLDSTENLYFQGIDPFTGCGGGGSGGGGSGGGGSGSRGHMPPL
TSAQQALTGTINSSMQAVQAAQATLDDFETLPPLGQDAASKAWRKNKMDESKHEIHSQVD
AITAGTASVVNLTAGDPAETDYTAVGSAVTTISSNLTEMSRGVKLLAALLEDEGGNGRPLLQ
AAKGLAGAVSELLRSAQPASAEPRQNLLQAAGNVGQASGELLQQIGESDTDPHFQDVLMQ
LANAVASAAAALVLKAKSVAQRTEDSGLQTQVIAAATQSALSTSQLVASTKVVAPTISSPVS
QEQLVEAGRLVAKAVEGSVSASQAATEDGQLLRGVGAAATAVTQALNELLQHVKAHATGA
GPAGRYDQATDTILTVTENIFSSMGDAGEMVRQARILAQATSDLVNAIKADAEGESDLENSR
KLLSAAKILADATAKMVEAAKGAAAHPDSEEQQQRLREAAEGLRMATNAAAQNAIKKGTGG
GGSGGGGSGGGGSC

Red = adaptions; Grey = expression tagged removed in production; C = cysteine attachment sites for attachment of the crosslinker.

**Protein expression and purification:** pGEL, GFP-VD1, mutated_R1, mutated_R2, and GFP were produced as codon optimized synthetic genes in pET151 plasmids (GeneArt), and transformed into BL21(DE3) cells. Overnight cultures were used to inoculate Lysogeny

Broth (pH 7.2) or M9 minimal media with $^{15}$N-ammonium chloride for labelled samples, and grown at 37°C until an $OD_{600}$ 0.6-0.8 was reached. The culture was then induced with 100 μM IPTG and grown overnight at 20°C. Harvested cells were resuspended in nickel Buffer A (20 mM Tris pH 8.0, 500 mM NaCl, 20 mM imidazole), sonicated and centrifuged (6,000 RPM, 10 minutes, 4°C) . The supernatant was then loaded onto a HP HisTrap Nickel column (Cytiva) connected to an AKTA start system (Cytiva) and eluted with nickel buffer B (20 mM Tris pH 8.0, 500 mM NaCl, 500 mM imidazole). Resulting pure protein was dialysed (10 kDa MWCO) into phosphate buffer (pH 7.4) overnight, ready for use. Labelled samples were TEV cleaved following HisTrap Nickel column and dialysed into Q buffer A (20 mM Tris pH 8.0, 50 mM NaCl). The resulting dialysed sample was loaded onto a HiTrap Q HP column connected to an AKTA start system (Cytiva) and eluted with Q buffer B (20 mM Tris pH 8.0, 1M NaCl). Resulting pure protein was dialysed into phosphate buffer (20 mM $NaH_2PO_4.2H_2O$, 50 mM NaCl, pH 7.4) overnight, ready for use.

**Circular Dichroism:** All circular dichroism (CD) experiments were performed on the JASCO J-175 spectropolarimeter using a 1 mm pathlength quartz cuvette. Far UV-spectra were obtained between 200-260 nm with an average of 4 scans at 100 nm/min, 0.5 nm step resolution, 1.0 second response and 0.5 nm bandwidth. For temperature scans CD wavelength was set to 222 nm. Measurements were taken between 20-90 °C with 20 second step resolution, 4 seconds of response and 1.0 nm bandwidth. Samples were prepared between 20-50 μM in 400 μL of phosphate buffer (20 mM $NaH_2PO_4.2H_2O$, 50 mM NaCl, pH 7.4).

**Protein NMR experiments:** Protein NMR experiments were conducted at 298 K using a Bruker AVANCE III spectrometer equipped with a QCI-P cryoprobe. $^{15}$N-labelled proteins were measured at 150 μM in 20 mM phosphate, 50 mM NaCl, 2 mM DTT and 5% v/v $^2H_2O$ at pH 6.5. Spectra were processed with TopSpin and CcpNmr Analysis 2.5.2.

**Gel Filtration:** Gel filtration was performed at room temperature using a Superdex 200 increase size-exclusion column (GE healthcare) system at a flow rate or 0.75 mL/min. Samples were run at 150 μM, or at 450 μM for the 3:1 condition, all at final volumes of 100 μL in 20 mM Tris pH 8.0, 150 mM NaCl, 2 mM DTT.

#### Section 2: Chemical structures

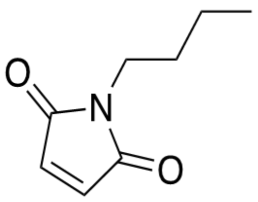

1

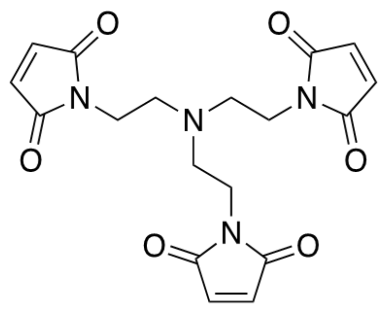

2

**Supplementary Fig. 1** Chemical structures of compounds **1** and **2**.

#### Section 3: Chemical synthesis and characterisation

##### Chemical synthesis:

**Crosslinker 1:** This compound was synthesised as described by Eloh *et al.*[1] with minor modifications. Maleic anhydride (1.00 g, 10.00 mmol) was dissolved in dichloromethane (DCM) (15.00 mL). *N*-butylamine (1.00 mL, 1.00 mmol) was added and the mixture was stirred at room temperature for 1 hour. The solvent was removed *in vacuo*, and the resulting white powder was re-dissolved in acetic anhydride (6.00 mL). To this solution, sodium acetate (0.50 g, 6.10 mmol) was added, and the mixture was heated at 80°C under reflux for 2 hours. The solution was diluted with distilled water (50.00 mL) and washed with diethyl ether (3 x 50.00 mL). The organic layer was collected and further washed with 0.1 M hydrochloric acid (1 x 50.00 mL) and 0.1 M sodium hydroxide (1 x 50.00 mL). The organic layer was dried over anhydrous sodium sulphate, filtered, and concentrated *in vacuo* to give the crude product as a colourless liquid. The N-butylmaleimide was further purified using silica chromatography, 85:15 (Ethyl acetate:hexane), producing a yellow oil with a yield of 11% (0.17 g, 11.00 mM). $^1$H NMR (400 MHz, 298 K, DMSO-$d_6$): δ: 7.01 (s, 2H), 3.38 (t, $J$ = 7.06 Hz, 2H), 1.46 (m, 2H), 1.21 (m, 2H), 0.86 (t, $J$ = 7.36 Hz, 3H) data was consistent with previously reported values.

**Crosslinker 2:** This compound was synthesised as described by Hanlon *et al.*[2] with minor modifications. A solution of maleic anhydride (0.59 g, 6.00 mmol) in anhydrous Dimethyl formamide (DMF) (2.43 mL) was prepared under inert atmosphere and cooled to 0°C. Separately, a solution of tris(2-aminoethyl)amine (0.29 mL) in dry DMF (2.04 mL) was prepared under inert atmosphere, and added dropwise to the maleic anhydride solution at 0°C over 30 minutes. The solution was stirred for a further 30 minutes at 0°C. A solution of sodium acetate (0.048 g, 0.60 mmol) in acetic anhydride (0.60 mL) was added to the reaction mixture at room temperature and stirred overnight at 50°C under inert atmosphere. The reaction mixture was concentrated using rotary evaporation, resuspended in DCM (50.00 mL) and washed with saturated brine (6 x 50.00 mL). The organic layer was collected, concentrated using rotary evaporation, resuspended in DCM (50.00 mL) and further washed with saturated sodium bicarbonate solution (6 x 50.00 mL). The organic layer was collected and concentrated using rotary evaporation to obtain the crude product. The crude product was purified using silica chromatography, 85:15 (Ethyl acetate:hexane). The resulting pure yellow crystalline product was dried under vacuum overnight with a yield of 9% (0.21 g, 0.54 mM). $^1$H NMR (400 MHz, 298 K, DMSO-$d_6$): δ: 6.98 (s, 6H), 3.38 (t, $J$ = 6.60 Hz, 6H), 2.60 (t, $J$ = 6.62 Hz, 6H) data was consistent with previously reported values.

#### NMR characterisation:

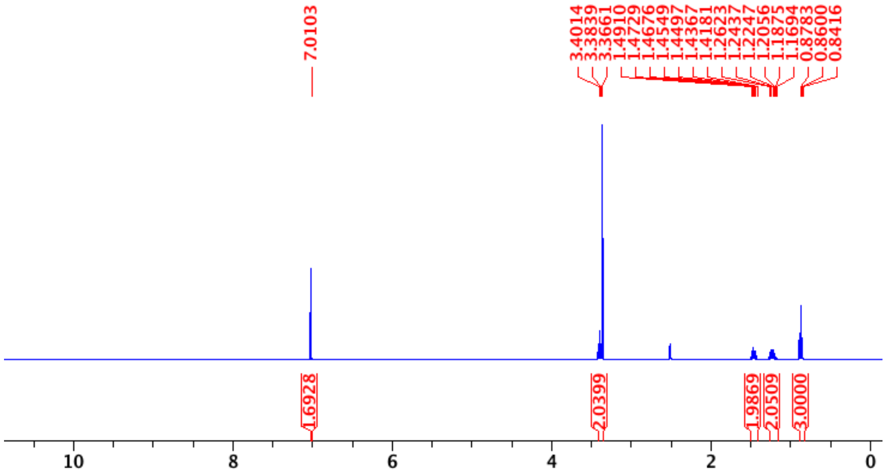


**Supplementary Fig. 2** $^{1}$H NMR spectra of crosslinker **1** in DMSO-$d_6$ conducted at 298.15 K.

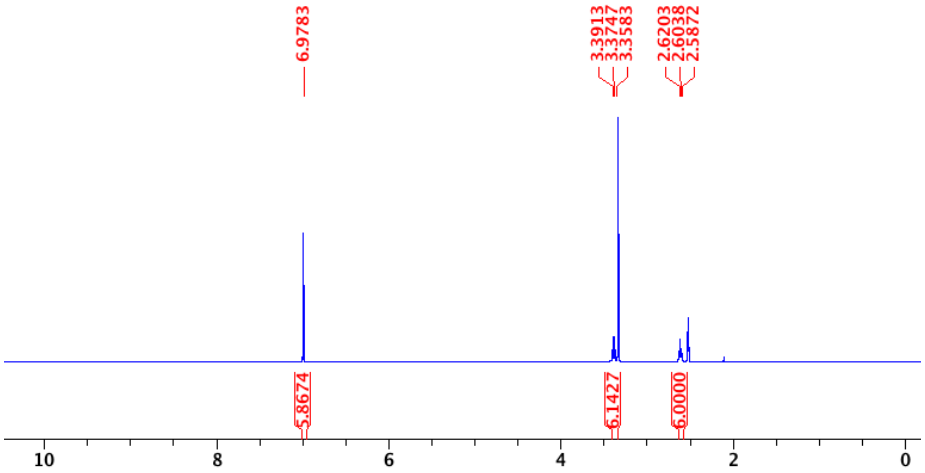


**Supplementary Fig. 3** $^{1}$H NMR spectra of crosslinker **2** in DMSO-$d_6$ conducted at 298.15 K.

#### Electrospray ionisation mass spectrometry of crosslinkers **1** and **2** forming substituted species with Kank1 KN-domain peptides.

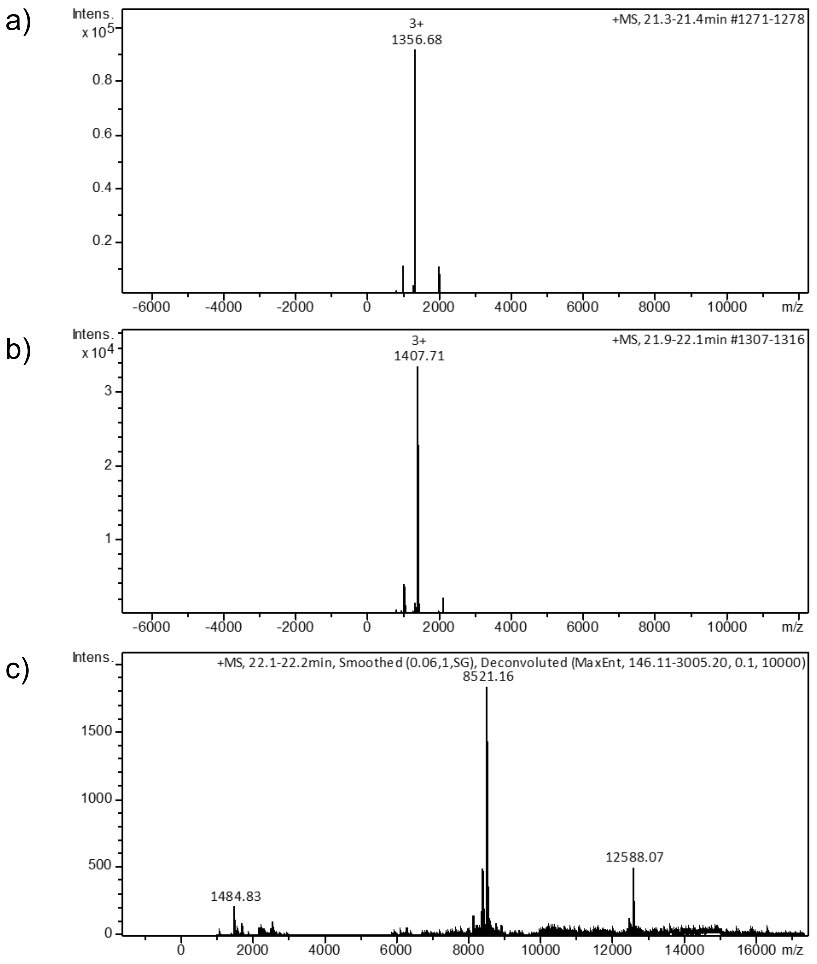


**Supplementary Fig. 4** LC-MS characterisation of crosslinkers **1** and **2**. **a**. Kank1 peptide (95% purity) (0.10 mM, 4067.01 g/mol) used for crosslinking characterisation. **b**. Compound **1** (1.00 mM, 153.08 g/mol) bound to a single Kank1 peptide (0.10 mM) confirming the maleimide group is capable of binding biological macromolecules. **c**. Compound **2** (1.00 mM, 386.12 g/mol) bound to three Kank1 peptides (0.10 mM) confirming all three maleimide groups are capable of binding biological macromolecules.

#### Section 4: pGEL characterisation

##### Circular Dichroism confirming preserved folding between pGEL and wild type R1-R3.

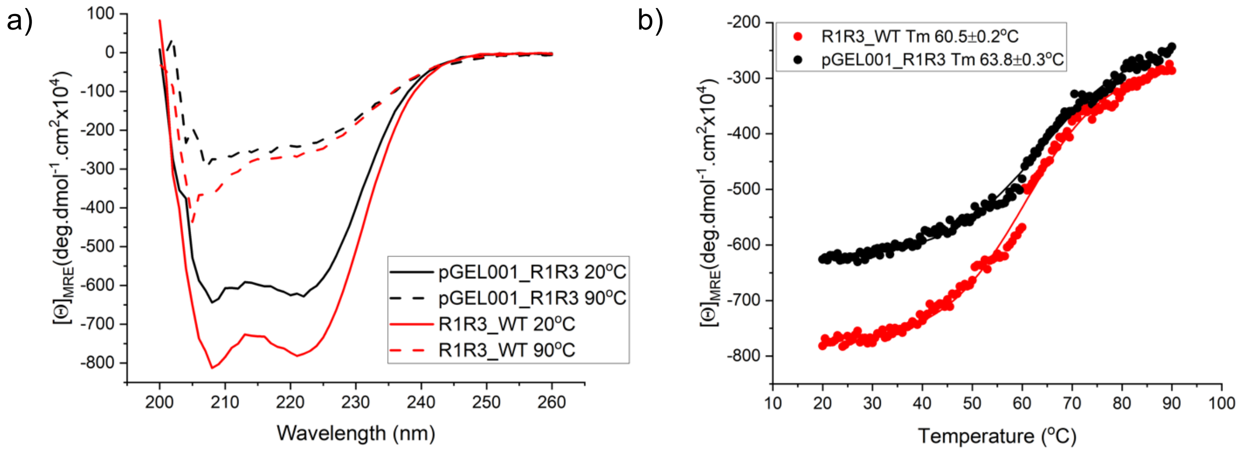


**Supplementary Fig. 5** Circular Dichroism analysis of pGEL and wildtype WT-R1R3. **a**. The CD spectra confirm that alpha helical folding is retained in pGEL similar to the wild type R1R3 following the cysteine-serine mutations (1 in R1 and 4 in R2). **b**. The thermal melt curves of the two confirm that only small changes in $T_m$ are observed between the two proteins.

##### Circular Dichroism confirming preserved folding between wild type R2 and the isolated mutated R2 domain from pGEL.

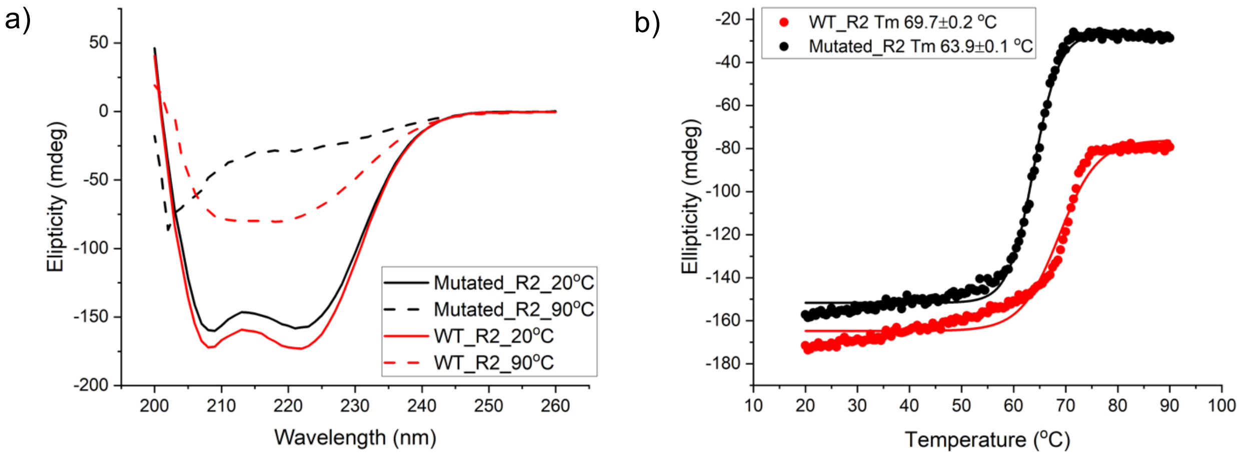


**Supplementary Fig. 6** Circular Dichroism showing **a.** Alpha helical folding is retained in the mutated R2 following four cysteine residues are mutated to serine. **b**. A change in $T_m$ is seen between the mutated R2 and wild type R2, suggesting minor alterations to stability has occurred.

2D HSQC spectra of $^{15}$N-labelled mutated R1 showing folding is retained compared to the wild type R1.

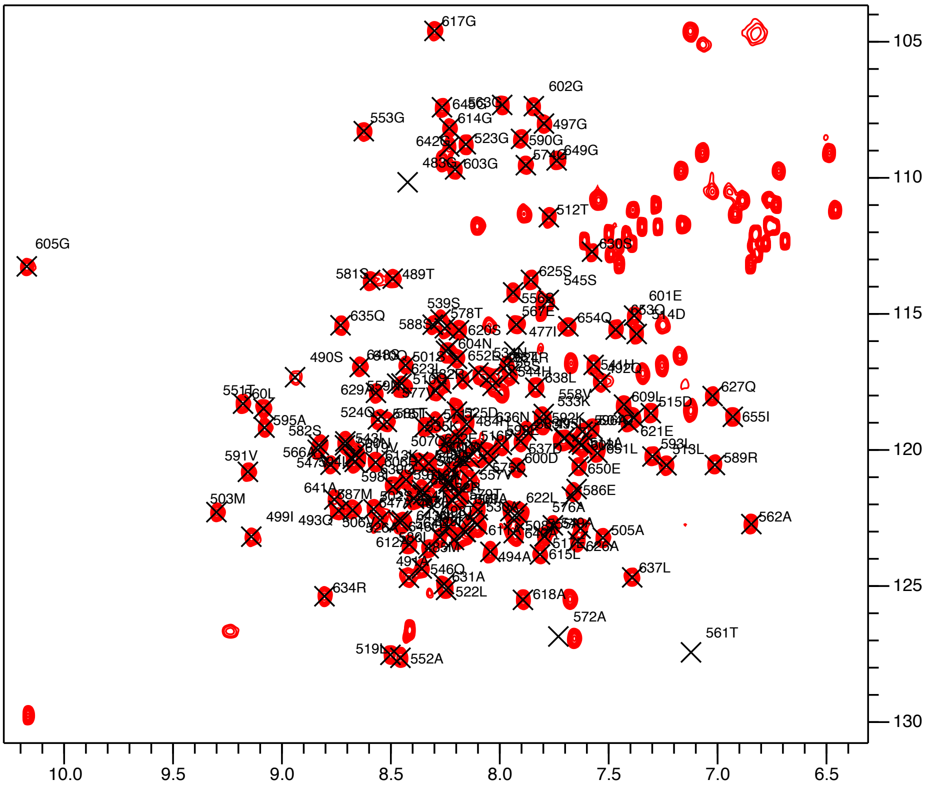


**Supplementary Fig. 7** HSQC spectra of $^{15}$N-labelled mutated R1 with wild type R1 assignments from Banno *et al.*[3] overlaid, showing that the single mutation in the mutated R1 does not perturb the folding of the domain.

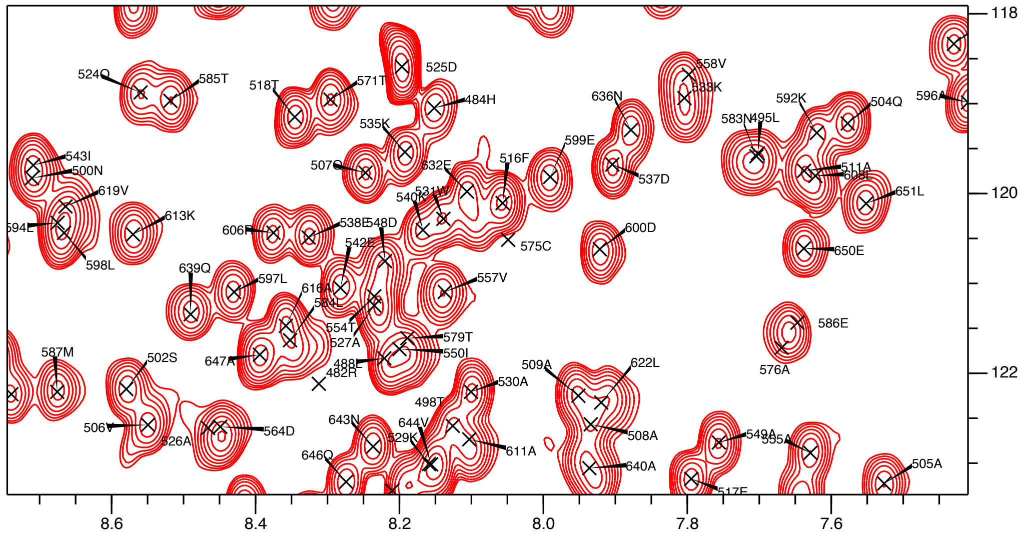

**Supplementary Fig. 8** HSQC spectra of $^{15}$N-labelled mutated R1 with wild type R1 assignments from Banno *et al.*[3] overlaid, centred on the region where the cysteine in wild type R1 is positioned.

### Section 6: Rheology of TSAM

#### Amplitude Sweep 1

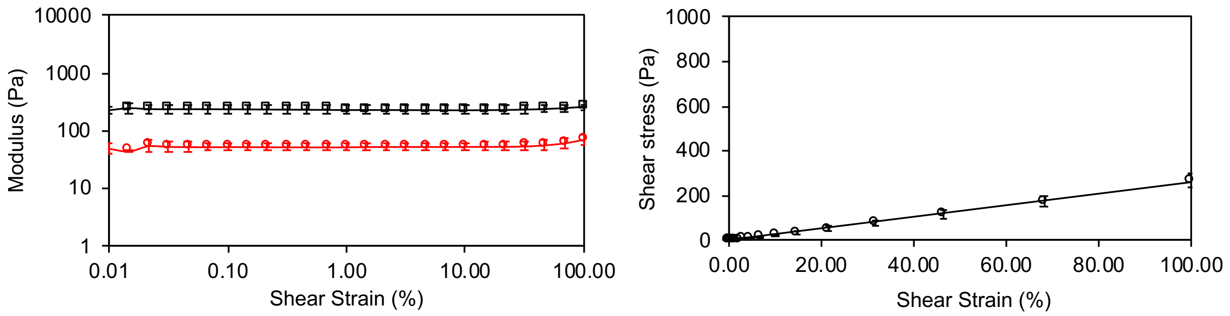


**Supplementary Fig. 9** Amplitude sweep 1 on TSAM (n = 3, error bars = SEM). (left) G' (black) and G" (red) against shear strain and (right) shear stress against shear strain.

#### Amplitude Sweep 2

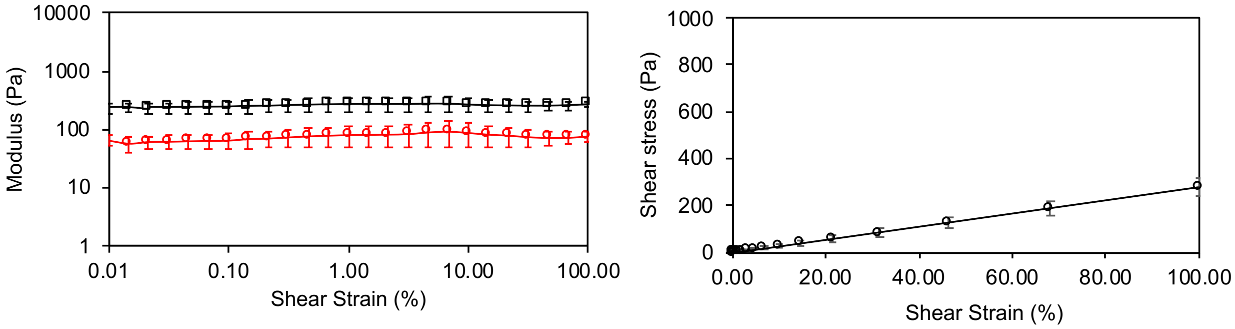


**Supplementary Fig. 10** Amplitude sweep 2 on TSAM (n = 3, error bars = SEM). (left) G' (black) and G" (red) against shear strain and (right) shear stress against shear strain.

#### Amplitude Sweep 3

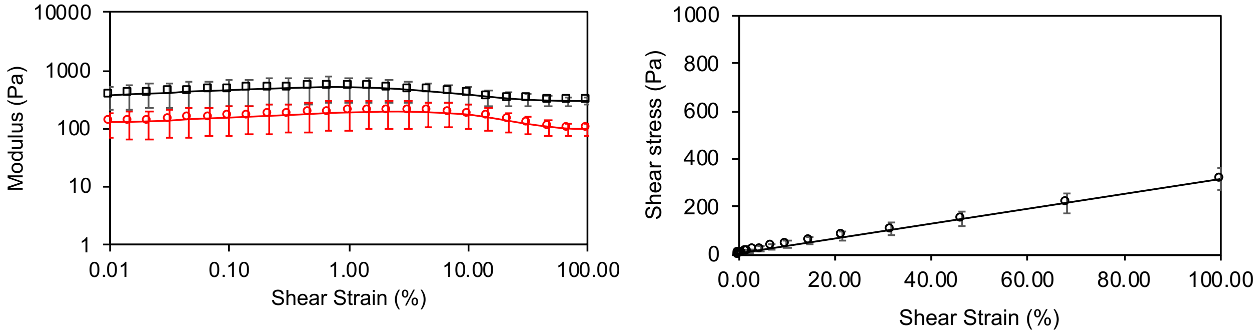


**Supplementary Fig. 11** Amplitude sweep 3 on TSAM (n = 3, error bars = SEM). (left) G' (black) and G" (red) against shear strain and (right) shear stress against shear strain.

#### Amplitude Sweep 4

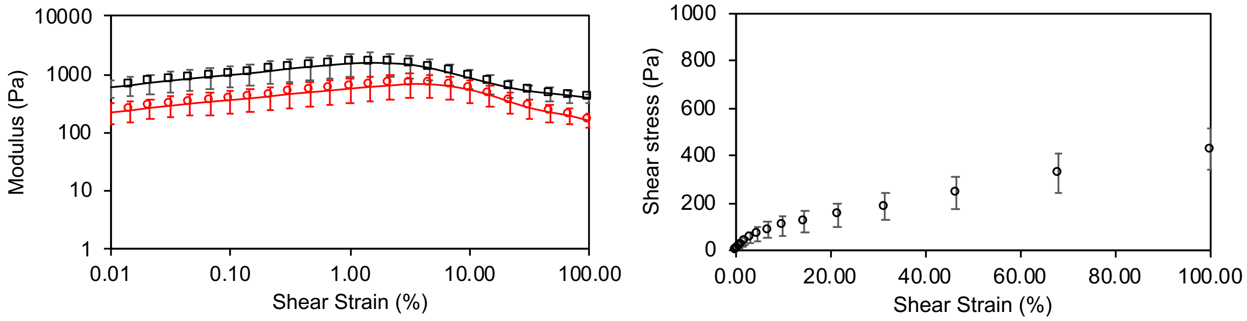


**Supplementary Fig. 12** Amplitude sweep 4 on TSAM (n = 3, error bars = SEM). (left) G' (black) and G" (red) against shear strain and (right) shear stress against shear strain.

#### Amplitude Sweep 5

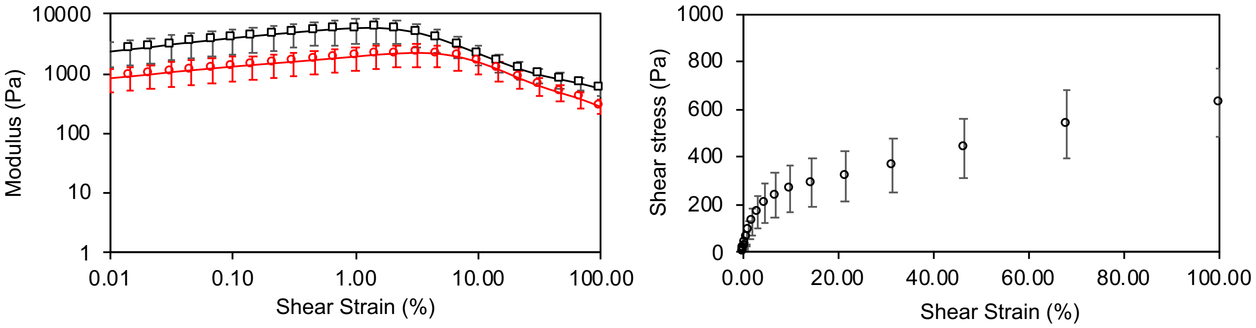


**Supplementary Fig. 13** Amplitude sweep 5 on TSAM (n = 3, error bars = SEM). (left) G' (black) and G" (red) against shear strain and (right) shear stress against shear strain.

#### Section 7: Analysis of TSAM with GFP-VD1, GFP and buffer

##### Gel filtration confirming GFP-VD1 binds to pGEL.

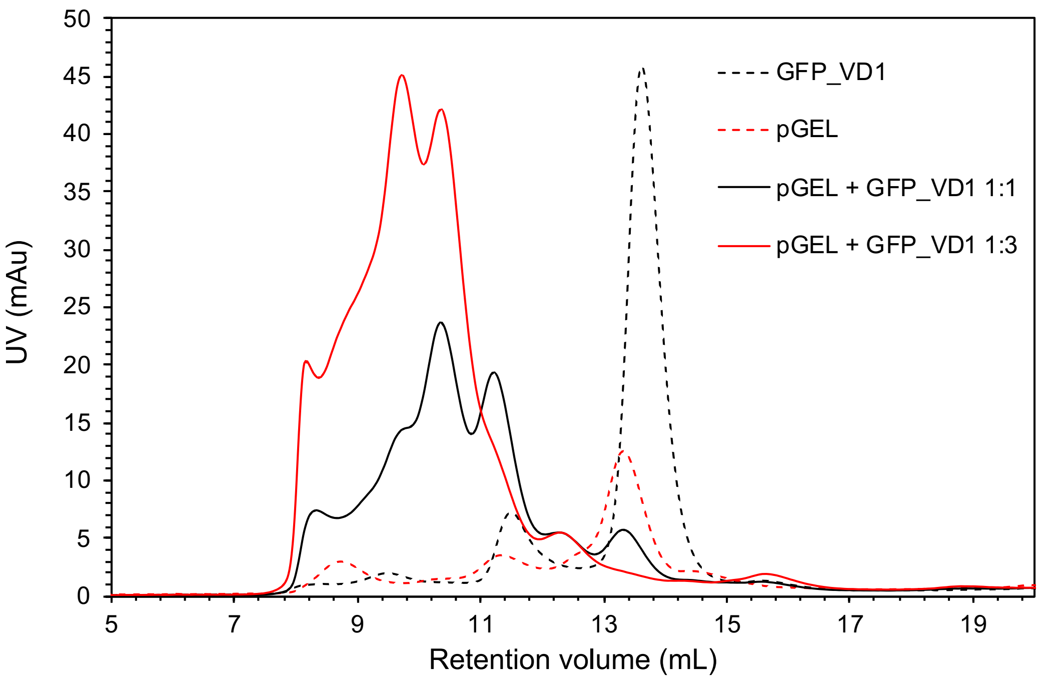


**Supplementary Fig. 14** Gel filtration profiles confirming GFP-VD1 binds to pGEL. GFP-VD1 = dotted black line, pGEL = dotted red line, GFP-VD1 + pGEL at 1:1 = solid black line, GFP-VD1 + pGEL at 3:1 = solid red line.

##### Amplitude sweep 1

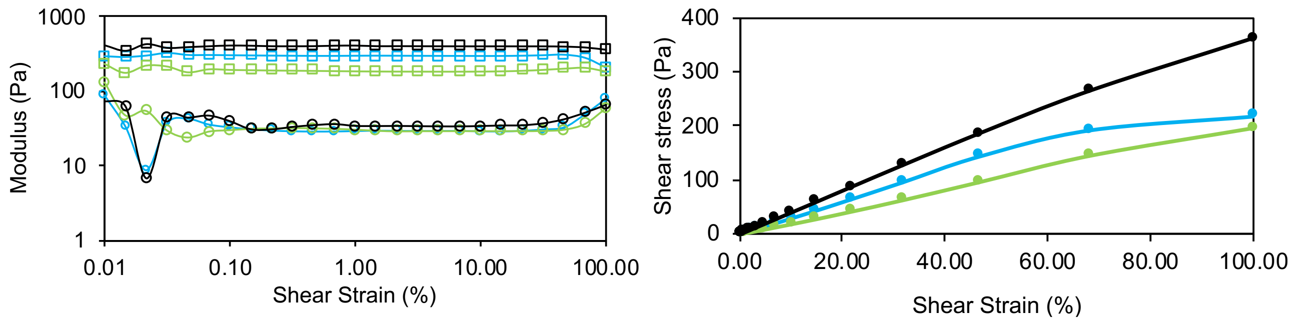


**Supplementary Fig. 15** Amplitude sweeps from rheological characterisation of TSAM after treatment with phosphate buffer (black), GFP (green) or GFP-VD1 (blue). G' and G" against shear strain and shear stress against shear strain for sweep 1.

### Section 8: Light gas gun (LGG) experiments

#### Supporting images from LGG experiments

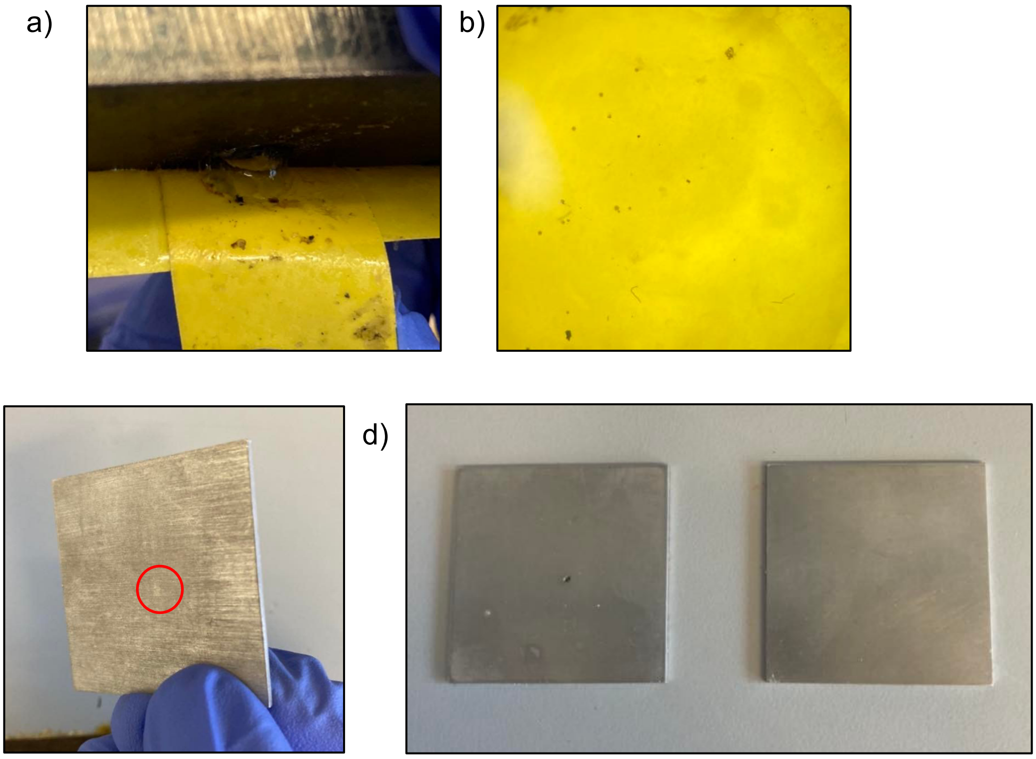


**Supplementary Fig. 16** Supporting images from light gas gun (LGG) experiment. **a**. Intact TSAM attached to tape on the back of the BTEA. **b**. Image under light microscope of basalt particles integrated into the TSAM shown in a. **c**. Resulting dent on the back of the aluminium back plate from the control LGG shot. **d**. side by side comparison of the back plates from the control LGG shot (left) and the TSAM LGG shot (right).

#### EDX analysis of caught basalt particles from light gas gun experiments imaged in SEM.

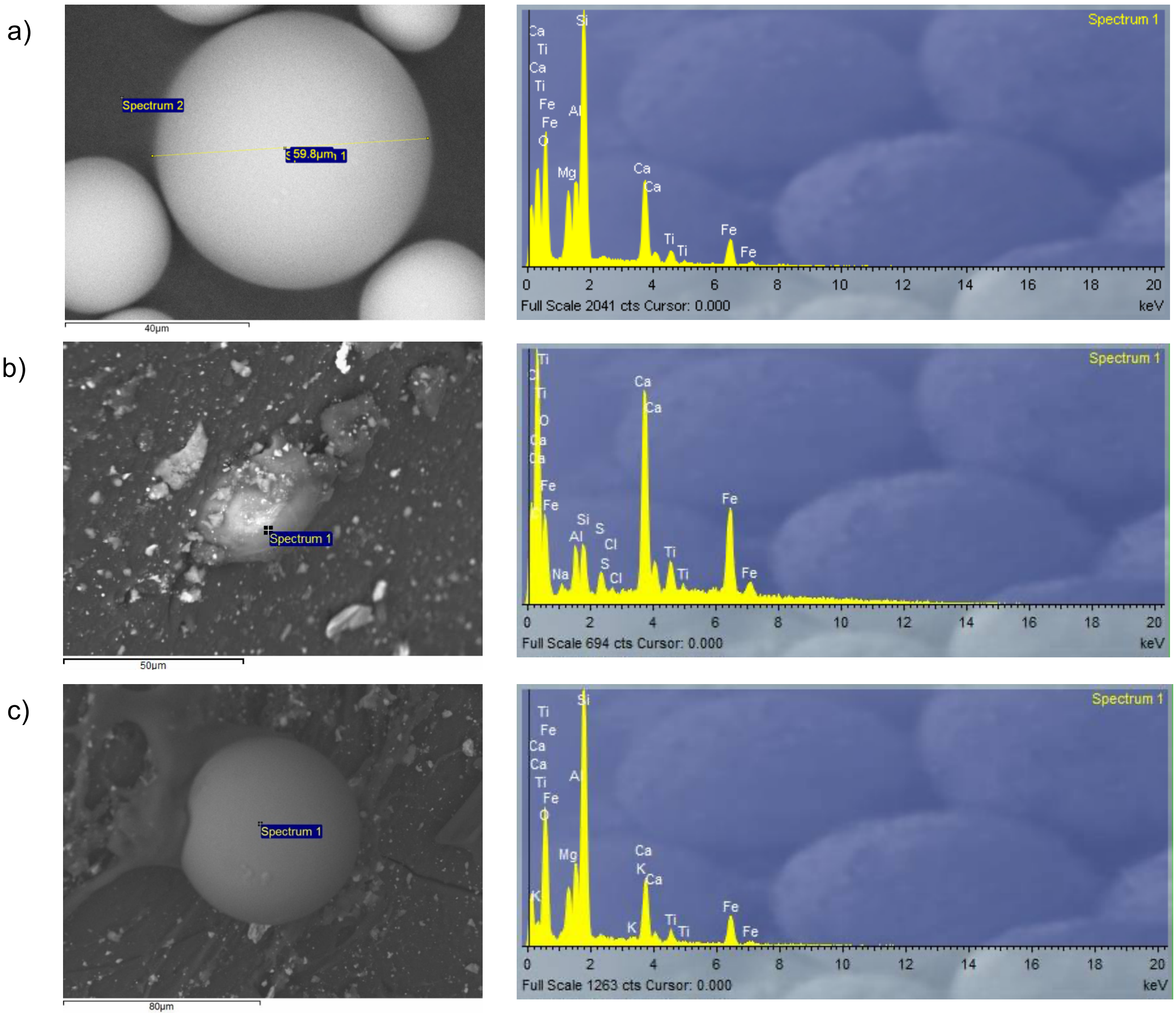


**Supplementary Fig. 17** SEM images of basalt particles and corresponding elemental dispersive X-ray (EDX) analysis. **a**. Basalt particle before being shot from LGG and its corresponding EDX analysis. **b**. Basalt particle 1 and its corresponding EDX analysis. **c**. Basalt particle 2 and its corresponding EDX analysis.

#### EDX analysis of caught burst disc shrapnel in light gas gun experiment imaged with SEM.

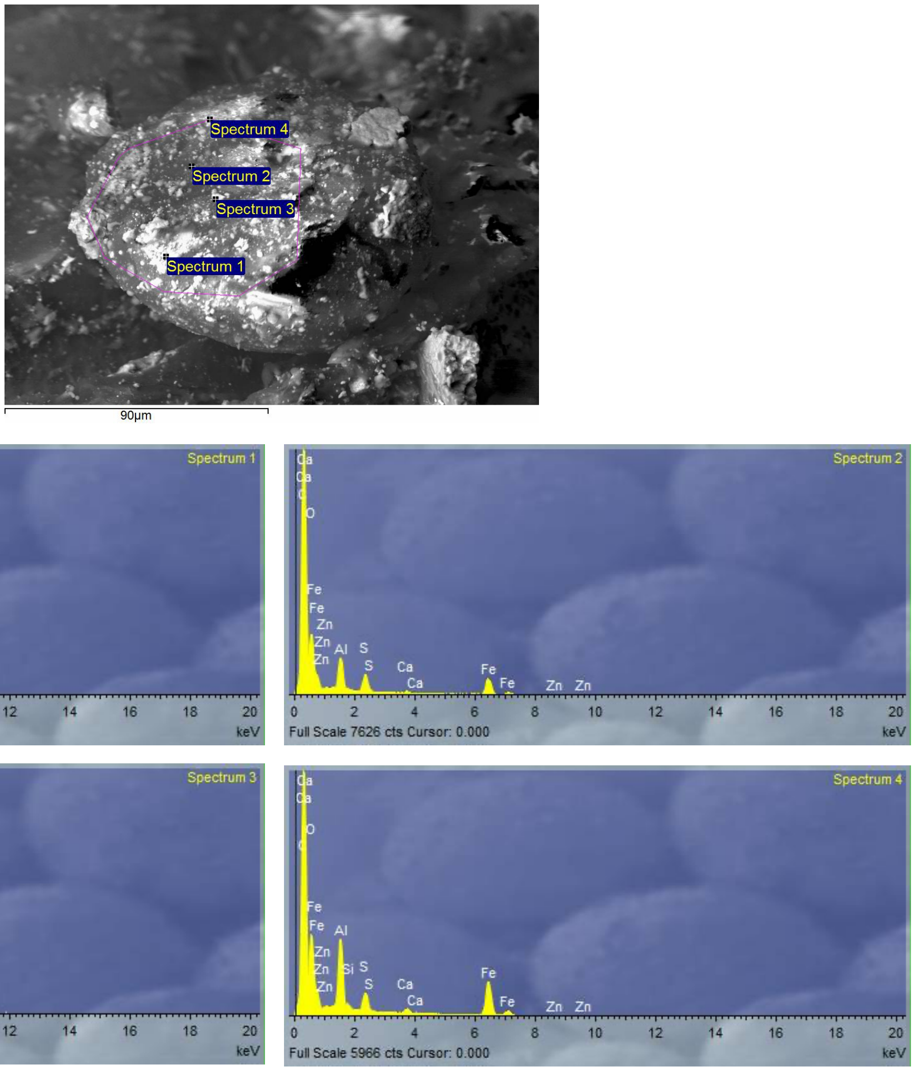


**Supplementary Fig. 18** SEM images of burst disk aluminium fragment and corresponding elemental dispersive X-ray analysis confirming aluminium.
